# ME-VAE: Multi-Encoder Variational AutoEncoder for Controlling Multiple Transformational Features in Single Cell Image Analysis

**DOI:** 10.1101/2021.04.22.441005

**Authors:** Luke Ternes, Mark Dane, Sean Gross, Marilyne Labrie, Gordon Mills, Joe Gray, Laura Heiser, Young Hwan Chang

## Abstract

Image-based cell phenotyping relies on quantitative measurements as encoded representations of cells; however, defining suitable representations that capture complex imaging features is challenged by the lack of robust methods to segment cells, identify subcellular compartments, and extract relevant features. Variational autoencoder (VAE) approaches produce encouraging results by mapping an image to a representative descriptor, and outperform classical hand-crafted features for morphology, intensity, and texture at differentiating data. Although VAEs show promising results for capturing morphological and organizational features in tissue, single cell image analyses based on VAEs often fail to identify biologically informative features due to uninformative technical variation. Herein, we propose a multi-encoder VAE (ME-VAE) in single cell image analysis using transformed images as a self-supervised signal to extract transform-invariant biologically meaningful features, including emergent features not obvious from prior knowledge. We show that the proposed architecture improves analysis by making distinct cell populations more separable compared to traditional VAEs and intensity measurements by enhancing phenotypic differences between cells and by improving correlations to other analytic modalities.

## Introduction

Understanding cellular changes and phenotypic pathways at the single cell level is becoming increasingly common as it allows for a more comprehensive understanding of cell state and cell-to-cell heterogeneity. Multiple analytical tools are available to extract, quantify, normalize, and evaluate single cell RNA sequencing (scRNAseq) data.^1,2,3^ However, until recently, analyzing single cell imaging data in a similar fashion was limited to extracting mean intensity profiles, predefined shape, textural and morphological features, and images stained with only a few markers. Emerging multiplexed imaging technologies such as cyclic immunofluorence (CYCIF),^4^ multiplexed immunohistochemistry (mIHC),^5^ CO-Detection by indEXing (CODEX),^6^ and Multiplexed ion beam imaging (MIBI)^7^ create images comprised of a large number of markers, expanding the depth of information captured. Robust analytical methods for high dimensional multiplexed imaging data, however, are still needed. One major limitation with analyzing highly multiplexed single cell images is the ability to extract biologically meaningful information on staining localization patterns that indicate divergent cell states. This problem is further confounded when relevant or novel features are obscured by non-informative features that can dominate a dataset. Imaging data has morpho-spatial information that is not captured using a simple mean intensity information of each marker, with successful quantification of these features potetntially leading to improved analysis and understanding.^8^

The classical approach for image feature extraction is to manually create a list of desired features and design a metric or algorithm that quantifies those features within the image. This traditional method is biased toward known and easily measured features and can miss subtle but important features. More robust image feature extraction has been employed using deep learning architectures such as the Variational Autoencoder (VAE)^9^ in other domains where feature representation can be automatically generated without supervising information or prior knowledge. However, the problem with VAE feature extraction in single cell imaging is that there are typically unimportant or uninformative features that drive differences between cells and skew the results in undesired ways.^10^ In single cell imaging data, these unimportant features include any form of basic image transformation such as rotation, transposition, affine/skew, and stretching. When these features are known and controllable transformations, they can be used for a self-supervised signal to remove the uninformative features and extract invariant features with respect to a set of transformations during model training. Despite having the same underlying information, the common uninformative features in transformed images distract deep learning architectures so that they ignore most of the biologically relevant features.^11,12,13,14,15^ This holds true in single cell images, where VAEs frequently ignore biologically meaningful features and focus on recreating the transformation features that have a high variance across the dataset. Whether in RNAseq or imaging data, untailored deep learning architectures are unable to overcome these uninformative features that skew analysis unless some modification is made to either their architecture or objective functions.^11,12,13,14,15^ Use cases of autoencoders for scRNAseq data have documented the difficulty of correcting for these features and extracting relevant information content into a single latent space. Many recent works propose changing autoencoder architectures to coupled networks or using multiple latent dimensions to overcome this without the need for biased hyperparameter tuning and data normalization, which are common practices in most scRNA analysis methods.^10,16,17,18^ Similar methodologies have also been explored in the imaging domain as well, seeking to correct transformative features with coupled networks, direct latent space modifications, novel layer architectures, and training networks with combinations of corrected and uncorrected image data.^11,19,13,20,12,14,15^ However, most of these corrected architectures are only targeted to one specified feature and are not able to generalize to other uninformative features without further modification.

Here we describe a novel method for single cell image feature extraction that removes specified uninformative features by making the features uniform and invariant across the reconstructions, using modified pairs of transformed input and output images by self-supervised transformation, and utilizing multiple encoding blocks. Using this multi-encoder VAE to control for multiple transformational features, we highlight its ability to extract more biologically meaningful and transform-invariant single cell information and better separate biologically distinct cell populations within a mixed dataset.

### Results

### A. Controlling for uninformative features

When a single transformational feature varies across a single cell imaging dataset, standard VAEs extract only the single dominant component to reconstruction. In a dataset where rotation varies from image to image, reconstructions along the principal component walk^21^ only constitute the angle of the cell and downstream analysis will be heavily skewed by this extracted component (Fig. 1a). In another dataset where polar orientation is the dominant feature, we observe the same behavior (Fig. 1b); VAEs only extract the dominant uninformative features and ignore many other subtle informative features necessary for detailed reconstruction.

**Figure 1:**
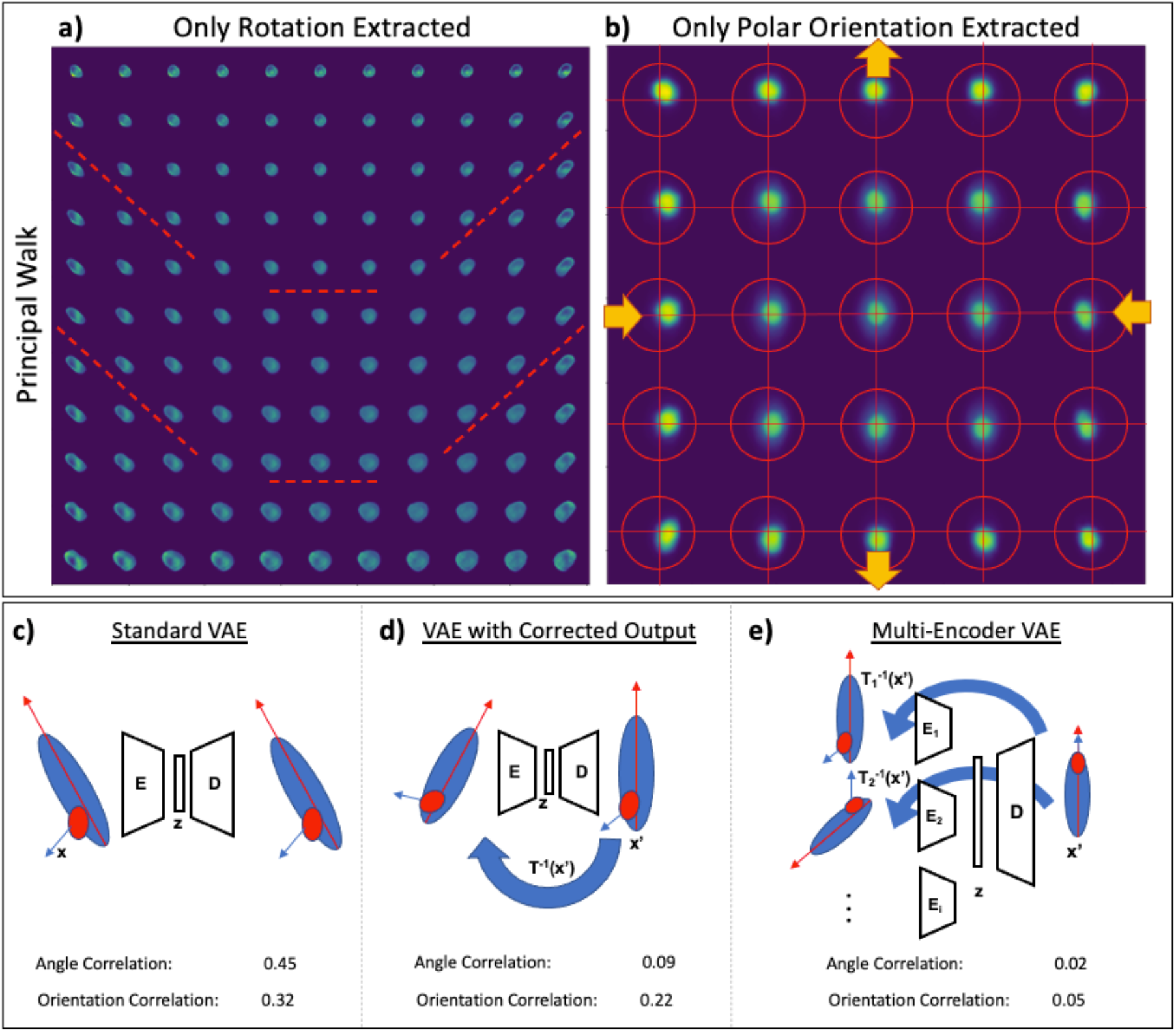
VAE hypersensitivity and proposed model architecture. VAE analysis of two datasets are shown, each governed by a single biologically uninformative feature **a)** rotation and **b)** polar orientation. Principal walk reconstructions^21^ show the VAEs’ governing features across the latent space through a range of image reconstructions. To correct this model hypersensitivity several architectures were tested: **c)** standard VAE with matched raw images. **d)** VAE with paired randomly transformed input and controlled output images **e)** the proposed multi-encoder VAE: VAE with corrections for multiple features (rotation, polar orientation, size, shape, etc.) using parallel encoder models, a shared latent space, and a single decoder model. In c)-e), a correlation between the embedding components and the respective feature (angle and orientation) is measured to quantify how effectively the model removes uninformative features.

In order to overcome model hypersensitivity to dominant uninformative features, several architectures were proposed and tested using transformed image pairs to make deep learning models learn the latent space by ignoring uninformative features (Fig. 1c,d,e). A standard VAE with no control for any uninformative features was used as baseline and shows a high correlation between the embedding components and the respective feature metrics (Fig. 1c and Supplemental Figure 1a). When a single factor is controlled (i.e. rotation), it becomes uncorrelated to all VAE encodings, and even the max correlated component in the latent space is insignificant (Fig. 1d and Supplemental Figure 1b). Controlling for one feature, however, does not significantly impact the other dominate transformation feature (i.e. polar orientation). With a double transformed output, attempting to correct for two features simultaneously, we saw decorrelation of both dominant features (Supplemental Figure 1c), but the reconstructed images of this architecture were poor (Supplemental Figure 1e) reflecting a failure of the model to learn any relevant feature embedding. The VAE with transformed output was shown to work on simple transforms that the model can learn such as rotation, but more complex features or pairs of transformations like rotation combined with polar orientation proved too difficult for the model. Finally, when both uninformative features were controlled for using the proposed ME-VAE with transformed image pairs, we saw a decorrelation in both uninformative features that were controlled for, indicating that the VAE reconstructions learned to overcome the multiple dominant features and focus on underlying features that better separate cell populations (Fig. 1e and Supplemental Figure 1d). Unlike with the corrected output VAE, the ME-VAE produced coherent reconstructions (Supplemental Figure 1f). Moreover, the proposed approach is both generalizable and scalable since it enables the control of many uninformative features together in a parallel combination by using a multi-encoder network where any number of encoders can be added in parallel and each encoder learns a single transformation.

### B. Improving Biological Interpretation on Single Channel Images

To evaluate the models’ ability to extract biologically relevant features with improved downstream usefulness, we analyzed a dataset (see Methods A) of single cell CYCIF images of MCF10A non-malignant breast epithelium cell line. The full dataset we analyzed is comprised of 6 ligand treated cell populations and is stained with 23 biomarkers. In this analysis, we restricted our analysis to populations that had been treated with PBS (control) and TGFβ+EGF ligands to induce a change in cell state. Analysis was done after 48 hours of exposure to ensure the most distinct results. We also limited the size of the images to only utilize the channel stained for Epidermal Growth Factor Receptor (EGFR). The EGFR marker was chosen for analysis because it displayed all uninformative features such as rotation, polarity, and cell size that we sought to correct. Furthermore, the two treatments were chosen because they have similar distributions of mean EGFR intensity and cell size, making them difficult to naively separate (Fig. 2a), but qualitatively show phenotypic differences such as compartment localization and stain texture. The proposed ME-VAE was tested to see if it could better separate and extract biologically distinct features between the PBS and TGFβ treated cell populations compared to a standard VAE within this labeled dataset. The separation was quantitatively measured by k-means cluster purity and qualitatively via separation in UMAP embedding space. Within this dataset we show that the ME-VAE is able to better separate distinct cell populations between the PBS and TGFβ treated MCF10A samples.

**Figure 2:**
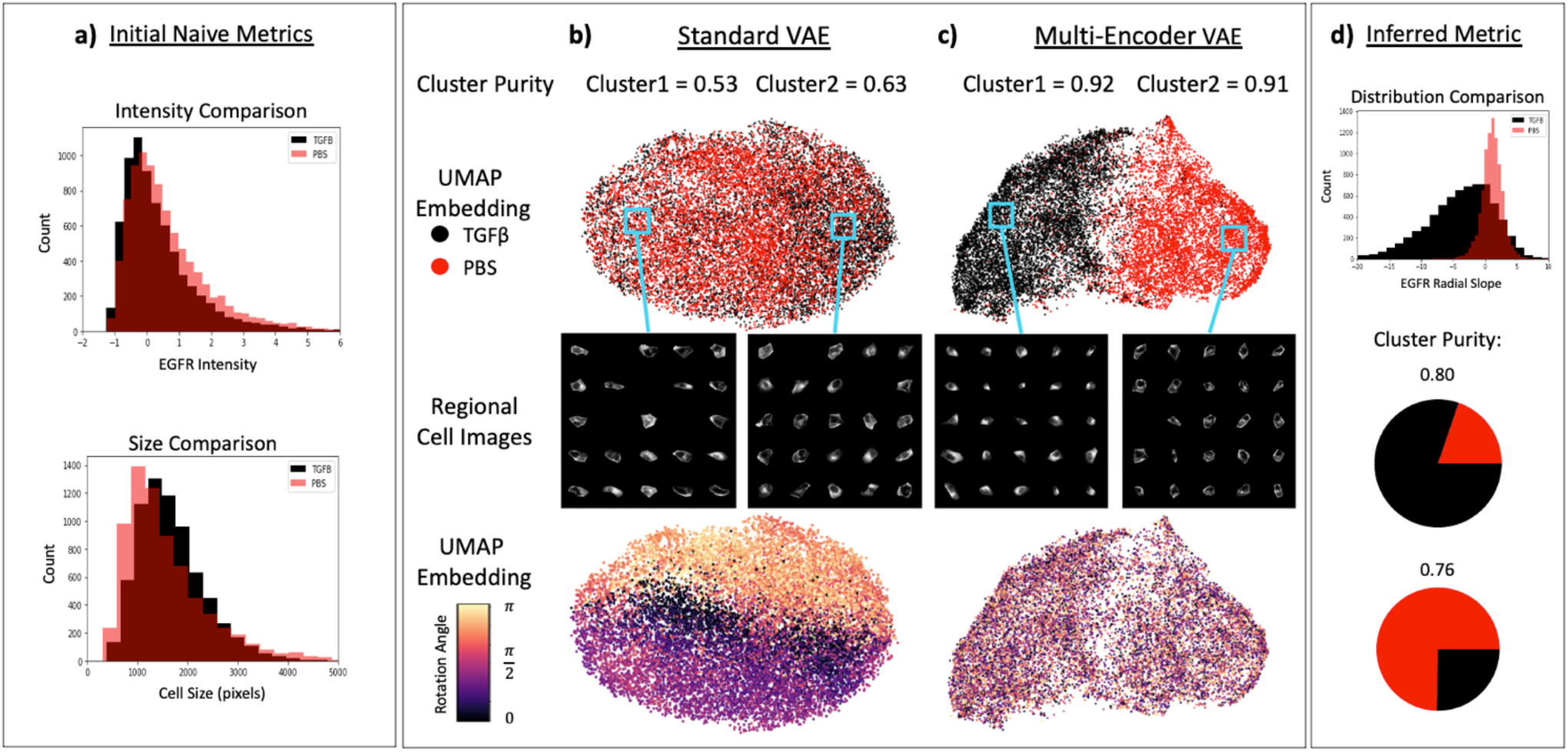
Separation of biologically distinct cell populations. **a)** Cells are compared using initial naïve metrics such as mean EGFR intensity and cell size to show the difficulty separating the cell populations. **b**,**c)** The models are quantitatively evaluated using cluster purity (k-means, n=2). Qualitative comparison is made using visual separation of two labeled cell populations in UMAP embedding space and visual analysis of cells from UMAP regions to identify biologically distinct factors. Rotation angle of cells are shown in UMAP embedding to show the influence of unimportant features on downstrean analysis. **d)** Cells are compared using radial slope (a metric inferred from visual analyzing the regional cell images in **c**).

As can be observed in Figure 2b, the standard VAE is incapable of separating the two cell populations, creating a mix of the labeled cell populations, both in k-means cluster space, as well as in UMAP embedding space. The cells within UMAP regions also have an arbitrary range of phenotypes, and the only patterns that can be observed are the patterns in uninformative features such as rotation. By comparison, the ME-VAE shows a dramatic increase in k-means cluster purity, and a clear separation of labeled cell populations in UMAP (Fig. 2c), indicating improved clusterizability and serperability of the data. All regional cell images within the multi-encoder’s UMAP space show distinct phenotypic differences that separate the cell populations with biologically relevant features (stain localization, intensity, and subcellular pattern). In PBS treated cell dominated regions, we see very dim EGFR staining throughout the cell with the stain being most heavily concentrated uniformly along the cellular membrane. TGFβ treated cells by comparison shows a cloudy diffuse concentration of EGFR stain throughout the cell with the heaviest concentration of stain localizing to one side of the nuclear membrane. These differences illustrate a clear difference in cellular regulation and compartmentalization of the EGFR protein induced by the TGFβ+EGF ligand combination. Several sub-clusters also form in UMAP space that subdivide the labeled populations, indicating features are being extracted that are able to identify different subpopulations within a treatment population, which can be used for more robust analysis. Based on observations from the multi-encoder output in Fig. 2c, we inferred that the metric of radial slope would better capture the biological differences between the two populations (see Methods C, Supplemental Figure 2). Using this metric, we see improved separation and cluster purity compared to the selected naïve metrics (Fig. 2a and 2d). The cluster purities from the inferred radial slope metric are still lower than the full ME-VAE cluster purity, indicating some other features are captured in the multi-encoder beyond the radial slope.

### C. Use Case with a Large Complex Dataset

The models were next trained on a larger portion of the CYCIF imaging dataset taken on the MCF10A breast epithelium cell line. The expanded set included treatment with five ligands and one control (PBS) for 48 hours and stained with the CYCIF marker set (see Methods A and Table 1). For this iteration, the ME-VAE was trained to control for cell shape and size in addition to the rotation and polar orientation performed previously as well as learned 23-channel images. This was done because the six treatments induced significantly different cell sizes that need to be controlled to improve the model’s ability to capture subcellular stain information.

In this larger dataset, the standard VAE performs similarly to the smaller dataset, encoding cells based primarily on dominant features such as size and rotation while largely ignoring complex staining information (Fig. 3a,b – left panel). Although visually there is some preferential localization (such as OSM on the left side and TGFβ on the right), it is clear that the ligand-treated populations are thoroughly mixed with poor separability. When looking at the intensity profiles, we can see that size has a strong impact on this left/right embedding space (Fig. 3b left). Most all other stains show little or no consistency within the embedding space, with the exception of DAPI and Ki67. These stains, however, show the same left/right distribution as size. It is likely that this distribution of staining intensities is simply a result of cell size since the whole cell mean intensity of a nuclear marker will decrease with larger cells and increase with smaller cells.

**Figure 3:**
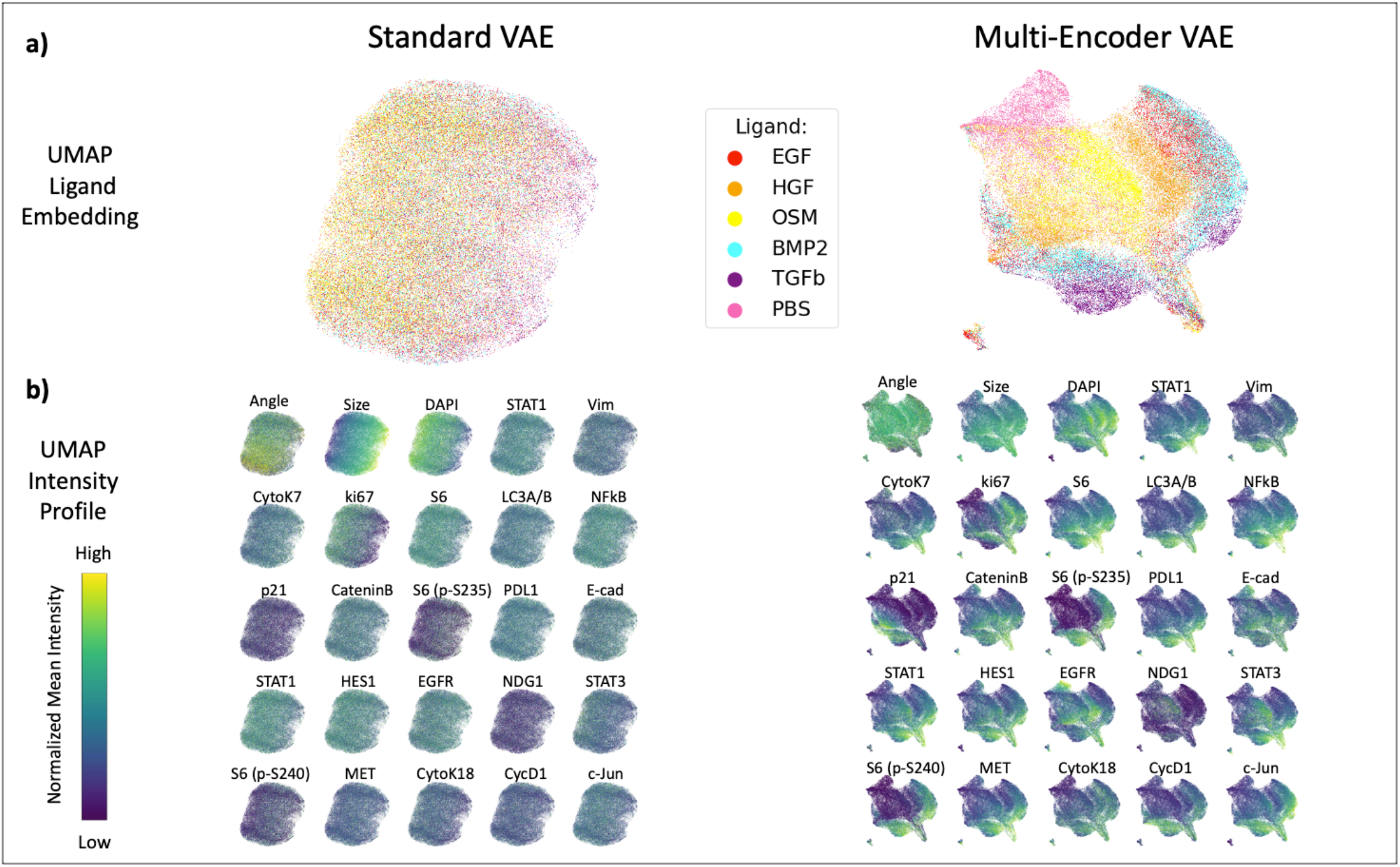
Ligand separation and feature distribution. **a)** UMAP embeddings for respective VAE encodings, allowing for qualitative visual evaluation of ligand separability. **b)** Distribution of stain features across UMAP space, colored by intensity.

Despite the increased complexity of the multi-channel CYCIF images and diversity of the dataset which could overload a simple architecture, the ME-VAE still shows good separation of the labeled cell populations (Fig. 3a - right). Subpopulations within a given label are even more distinct in this larger dataset, with multiple clusters for HGF, BMP2, and TGFβ. By analyzing intensity profiles and regional cell images of these subpopulations, we can see differences in expression (Fig. 3b right and Supplemental Figure 3b - right). The UMAP intensity profiles also show clear stain intensity patterns indicating that the ME-VAE encoding space contains relevant biological information. The distribution pattern of size does show some presence in the UMAP space, but the effect is largely dulled in comparison to the standard VAE.

There are many differences in stain expression that can be observed between and within the ligand-treated cell populations. Here we discuss some of the most noticeable drivers of separation. In comparison with all other treatment groups, PBS shows a marked deacrease in Ki67 expression. This is consistent with a relative decrease in proliferation compared to other ligand treatments. The TGFβ populations show an increase in S6 expression relative to other ligand populations, indicating an increase in cell growth. This can be observed visually by looking at the regional cell images (Supplemental Figure 3b right); however, it is worth noting that this high expression of S6 is seen in both large and small cells treated with TGFβ. In both EGF and BMP2 treated populations, decreased expression of membrane adhesion proteins such E cadherin and β-Catenin was observed. This decreased expression presents visually as dim stain, but the marker is still localized to the membrane and is not missing or diffuse throughout the cell.

More interestingly, there are also differences in stain distribution that can be observed through the model. In both TGFβ- and PBS-treated cells, we see increased concentration of HES1 localized primarily to the nucleous, while in other populations the distribution is even throughout the cell. In the case of TGFβ-treated cells, this localization is accompanied by a large increase in expression intensity, but in the case of PBS, it is not significantly more intense than some of the other ligand-treated populations. In a similar manner, Stat1a is observed to be primarily located in the nucleous for TGFβ-, BMP2-, and OSM-treated populations, but shows decentralized staining in cell images for other ligand populations. This is important because both HES1 and Stat1a are functional in the nucleus (Stat1 particularly as it translocates into the nucleous as part of its functional pathway) with limited activity in the cytosol.^22,23^

One final observation worth note is that p21 shows a unique ability to separate very distinct subpopulations in TGFβ-, HGF-, and BMP2-treated cells. This would indicate that within these ligand populations, there are subsets of the population that are undergoing growth arrest due to inhibition of cell cycle progression via p21 regulation.

It is clear from these biological observations that the ME-VAE is able to capture relevant biological information and separate cell populations in a way that highlights important features without significant interference from the unformative features that were controlled for. Furthermore, the ME-VAE can capture emergent biologically relevant imaging features not obvious without prior knowledge. By contrast, little to no biologically relevant information can be easily obtained from the latent encodings of the standard VAE.

### D. Correlation with Pathway activity profiles via Reverse Phase Protein Arrays (RPPA) and ME-VAE features using CYCIF

By reordering VAE features using hieachical clustering to form a feature spectra, we can extract broader patterns from the VAE and reduce the dimensionality of the feature set. While the standard VAE shows very poor self correlation, with only a handful of features clustering together with significant correlation (Supplemental Figure 4a top), we observe a very clear pattern of self-correlations between ME-VAE features indicating the model is successfully extracting several distinct expression patterns and that there are clear clusters of VAE features that encode for similar things (Fig. 4a top). Using hierarchical clustering, we identify ten representative clusters from the ME-VAE latent space that illustrate different expression patterns. We can visually observe the patterns using a representative image for each cluster (Fig. 4a bottom). Using these feature clusters, we can create aggregated feature sets using the mean of all features in a cluster. Representative cell images were chosen by selecting cells that had high expression of their respective cluster to understand and interpret VAE aggregated feature clusters. Between clusters 0 and 1, we can visually see a difference in the ratio of nuclear size and cell size. Cluster 1 encodes for larger nuclei than is encoded in cells from cluster 0 (this pattern is reaffirmed in Fig. 4b where cluster 1 is correlated to DNA pathways and CYCIF stains while cluster 0 does not). Cluster 4 is a highly varied cluster, but contains large cells with more diffuse intensity patterns. From these clusters of VAE features we can see that the ME-VAE architecture is extracting a combination of intensity and morpho-spatial profiles with at least 10 clear axes of variation. Using these reduced feature sets, we can analyze for biological meaning with fewer spurious correlations than comparing many to many.

**Figure 4:**
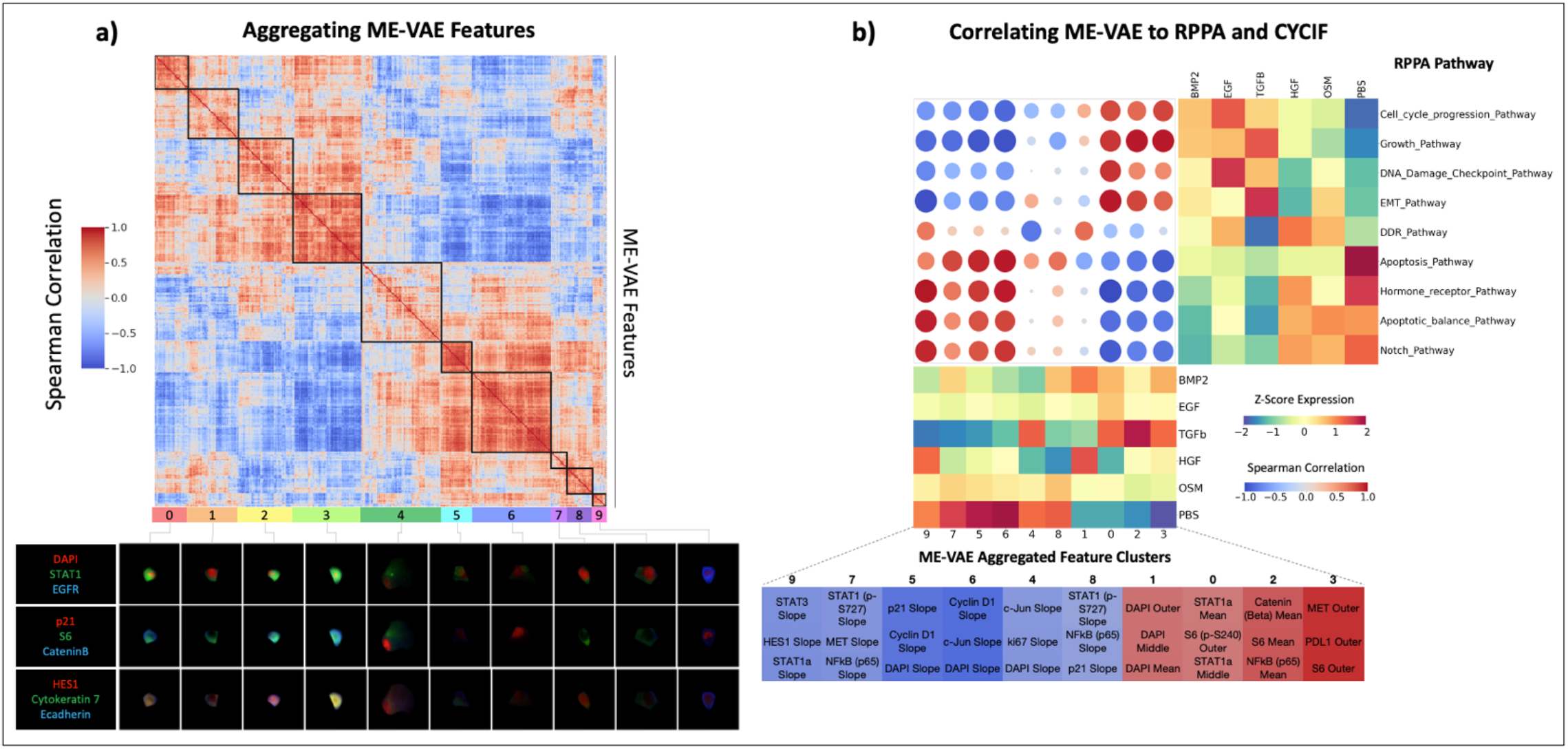
ME-VAE feature aggregation and transitive inter-modality correlation. **a)**Using the single cell observations as features, correlations are drawn between pairs of ME-VAE features. These features are then hierarchically clustered to observe patterns and reduce VAE features to aggregated feature sets. Cell images were assigned aggregated feature scores using the mean expression of each feature in a cluster. Shown are representative cells that are highly expressing for each respective cluster. **b)**Correlation matrix between RPPA pathway activity scores and ME-VAE aggregated features. Samples from the two modalities were paired by their ligand treatments, resulting in a sample size of n=6. RPPA pathways and VAE features were hierarchically clustered to show prominent patterns in correlation. ME-VAE aggregated features were also correlated to several metrics of CYCIF expression (mean inner, mean middle, whole cell mean, and radial slope) for all 23 stains. This CYCIF correlation was done using the full dataset of single cell images (sample size n=73,134). The table of CYCIF correlations shows the top three correlations for each ME-VAE aggregated feature.

A growing method for single cell analysis is to integrate multiple modalities. Multi-modal integration helps to validate where the two modalities overlap, expands the dataset with mutually exclusive or orthogonal features, and allows for cross-wise mapping of features. This is important for VAE based single cell image analysis because it frames inherently obscure encoding features in a biological context and validates that the extracted features are coherent. The increased feature range extracted by the ME-VAE allows for the cross-wise mapping and integration of complex CYCIF image features and other modalities, in this case reverse phase protein arrays (RPPA).

When correlating the 7 aggregated standard VAE features with RPPA pathway activity, we notice two very distinct issues. First, there is only a single standard VAE aggregated feature that shows any significant pattern of correlation, and that aggregated feature correlates to every RPPA pathway activity profile (Supplemental Figure 4b). The second is that there is a single RPPA pathway that correlates to every standard VAE aggregated feature. When the datapoints of the standard VAE are shuffled, we notice a similar overall magnitude of correlation values (data not shown), indicating that the correlations found between the standard VAE and RPPA pathways are likely spurious. When correlating standard VAE aggregated features to the extracted CYCIF metrics (Methods C, Supplemental Figure 2), the Spearman correlations are small despite the increased sample size of n=73,134 (Supplemental Figure 4b), with the largest correlations being restricted to nuclear markers such as CyclinD1, DAPI and Ki67. As mentioned above this is likely an artifact of encoding for size as nuclear expressions can be a function of cell size.

By contrast the ME-VAE features show good variation and distinct patterning between all ligand treatments (Fig. 4b, Supplemental Figure 4), resulting in more powerful and informative Spearman correlations with RPPA pathways. Unlike with the standard VAE, when the datapoints are shuffled, we see a significant decrease in correlation strength (data not shown), indicating that the observed correlations are less likely to be spurious. The improved correlations illustrate the multi-encoder’s better applicability for multi-modal integration and comparison. Correlations between ME-VAE aggregated features and CYCIF metrics were also performed using Spearman correlation using the full 71,314 sample size. For all 10 aggregated features, we see strong and consistent Spearman correlations. These correlations illustrate that the aggregated features extracted by the ME-VAE have biological interpretability in both CYCIF and RPPA and builds confidence in the model’s ability to capture biologically relevant information.

These biological correlations are validated by looking at representative images for each ligand treatment (Supplemental Figure 3 and 5). The stains shown were selected because they had high correlations to the aggregated ME-VAE features or showed distinct patterns visually. The same patterns observed in the CYCIF correlation table and VAE Z-score expression matrix (Fig. 4b), are also qualitatively confirmed by visual inspection. For example, S6 expression (ME-VAE feature 0) is high in BMP2, EGF, and TGFβ, and it is low in HGF, OSM, and PBS. Radial CyclinD1 radial slope (ME-VAE aggregated feature 6), as shown in Supplemental Figure 2, is negative in BMP2, EGF, and TGFβ, with high stain intensity in the inner compartment and rapid decrease toward the cell perimeter; conversely, HGF, OSM, and PBS show much dimmer CyclinD1 expression in the inner compartment. This pattern is even more clearly seen in the radial HES1 slope (Supplemental Figure 5), where HGF, OSM, and PBS show a more continuous stain abundance all the way to the cell membrane. Although the RPPA sample size (n=6) is still too small to achieve statistical significance, the correlations between protein markers in CYCIF and RPPA pathways linked by VAE features, are supported by known literature. We observe a correlation between DAPI expression (ME-VAE aggregated feature 1) and DNA damage and repair (DDR) pathway, which we interpret to be consistent with known biology. A more interesting example (ME-VAE aggregated feature 9) shows a strong correlation between the Stat3 radial slope of distribution and the epithelial-to-mesenchymal transition (EMT) pathway and hormone receptor pathway in RPPA. This is validated by prior literature showing that Stat3 distribution throughout the cell, its activation in the cytoplasm, and its translocation to the nucleus are important in the EGF induced epithelial-to-mesenchymal transition pathway.^24^ The ME-VAE architecture is also capable of extracting patterns when multiple markers play a role; an example of this is CyclinD1 and p21 (ME-VAE aggregated feature 5) which is known in literature to play a joint part in the cell growth pathway.^25^ These observations support the relevance of ME-VAE features and demonstrate a potential application of multi-modal integration by using the proposed approach for single cell image analysis.

The ME-VAE can also improve downstream analysis by increasing population separability (Fig. 5) as measured by mean pairwise Tukey p-values and mean effect sizes to represent the separability of feature distributions between ligand pairs. For the given MCF10A dataset, the CYCIF markers were chosen with the known ligands and cell populations in mind to highlight differences between the populations and separate them. This results in already decent separability using just CYCIF mean intensity information (Fig. 5 left). Within the CYCIF mean intensity features, the small mean Tukey pairwise p-value, indicates that for a given feature many of the populations are significantly different, and also have a decent effect size. Because ME-VAE features are able to capture more detail and nuance than than the selected naïve intensity metrics, the multi-encoder is able to improve the separability. ME-VAE features show lower mean Tukey pairwise p-values indicating a greater average significance in separability, and the effects sizes for those separations were larger (Fig. 5 right). The exception to this, shown in the first example, is S6, where the CYCIF mean intensity shows better separability, but even in this example the multi-encoder’s feature is still adequate. It is worth noting that ME-VAE latent space features will work in combination to represent even a single stain, and so separability can be expected to improve even further when utilizing more than just one feature at a time.

**Figure 5:**
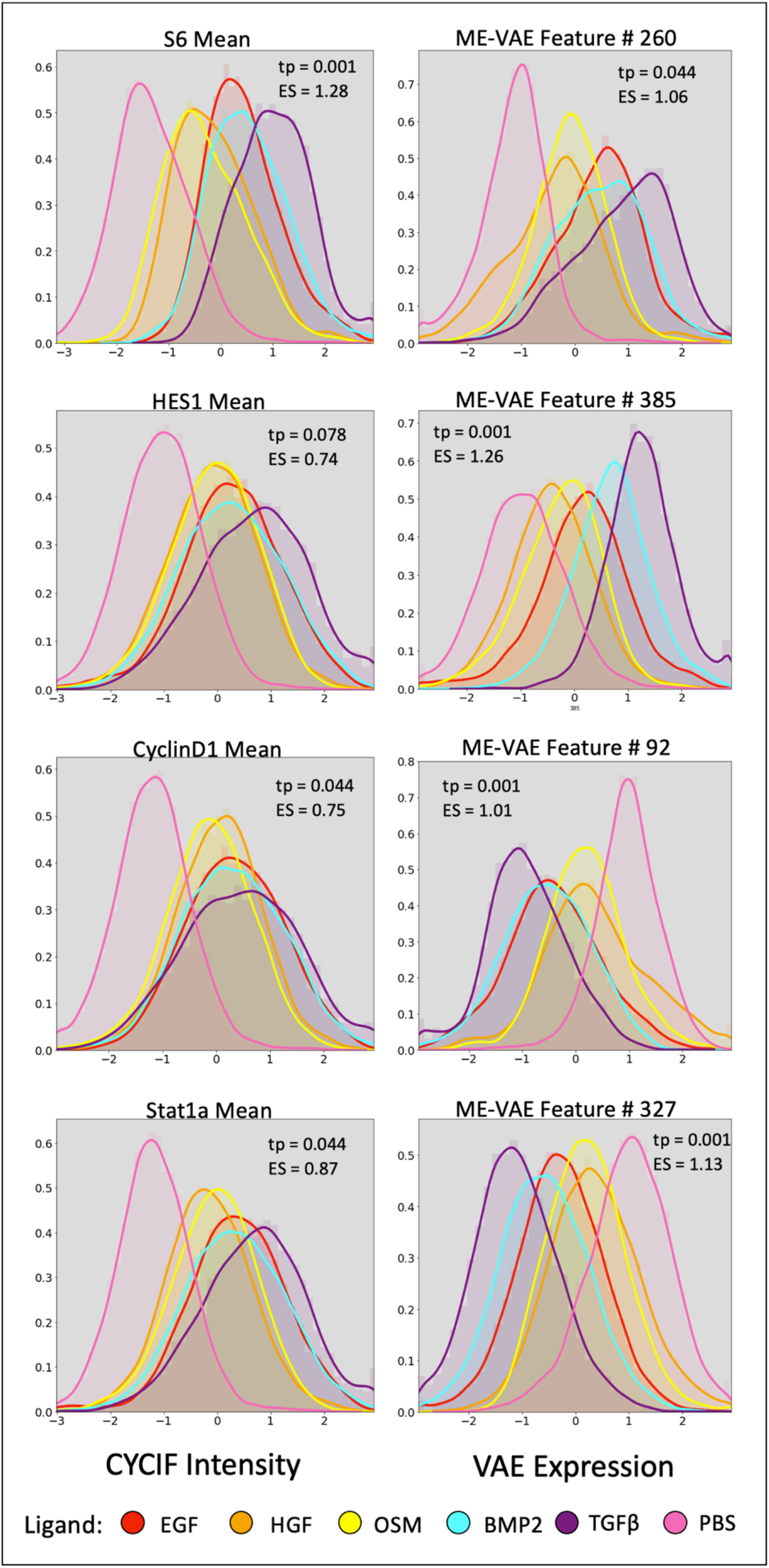
Separability of ligands. Density function and histogram for several CYCIF and ME-VAE feature pairs. The mean Tukey-pairwise p-value (tp) and mean effect size (ES) shown for each feature.

## Discussion

Just as it is necessary to pre-process, normalize, and remove unwanted features from scRNAseq or RPPA analysis, so too is it necessary to remove uninformative features from single cell imaging analysis in order to better extract features of interest. Without this guided feature alignment, VAE applications for single cell image analysis will only reconstruct dominant features while ignoring subtle more informative features (Fig. 1a, b). Transform invariant autoencoders for image analysis have been used before, but controlling for the multiple complex features present in single cell imaging data is difficult given a single encoder network. By making uninformative features invariable across a dataset, using pairs of transformed images in parallel encoding blocks (Fig. 1e), VAE priority can be shifted to mutually shared biologically relevant information (Fig. 2c, Fig. 3). This also results in a more complex and meaningful latent space, as more encoding features are necessary to represent the complex features being reconstructed.

Feature extraction is important for all downstream analysis and interpretation, but often times naïve metrics are not sufficient to capture biological differences and separate cell populations, especially in datasets where labeled populations are not known beforehand. By separating populations with the ME-VAE, distinct populations and biologically meaningful metrics can be established allowing identification of emergent properties from images such as distribution and staining pattern (Fig. 2c, Fig. 3, Supplemental Figure 3), with increased separability compared to naïve metrics using intensity information alone (Fig. 2c/d, Fig. 5). This is because a well-trained encoder can capture complex patterns such as staining texture, pattern, and distribution that are not intuitively quantified by other metrics. Although theoretically an infinite number of handcrafted naïve features could be crafted to capture more information, the advantage of deep-learning extracted features is that it can extract the most important features of an image without need for prior knowledge or feature selection. More complex single cell analysis methods such as multimodal integration and comparison (Fig. 4) require that a wide range of features are extracted that are reflected biologically. The ME-VAE architectyre provides an important step for biological research by linking imaging data to molecular readouts. By employing this architecture to extract a larger range of features and metrics from single cell images, potential applications, such as multi-modal integration using imaging features, become available which were previously restricted due to inadequate cell representations.

The simplicity of the multi-encoder design makes it easily incorporated into more complex deep-learning architectures, such as being augmented with a discriminator to improve reconstruction quality. This methodology is limited by the fact that the uninformative features being corrected for must meet two criteria: 1) being known beforehand so that it can be addressed with a new encoding block and transformed image pair; 2) be a known transform operation such as rotation, affine, or scale such that a respective randomly transformed image can be made using the operation. Despite this limitation, the majority of dominant uninformative features are based on known transformations, making the ME-VAE architecture widely applicable. Computationally the model is not significantly larger than a standard VAE or other comparable architectures, as it only adds a single encoding block for each undesired feature. The majority of increased computation time comes from the image transformation steps. Future applications of this architecture will allow complex features such as texture, pattern, and distribution to be extracted from single cell images free from the hassle of disentangling dominant uninteresting transform features. Images contain morpho-spatial features not shared by their other single cell counterparts (scRNAseq and RPPA), and by implementing this architecture, the scientific community will be able to analyze these unique image features with the same robustness as algorithms made for the other well-established single cell modalities.

## Methods

### A. Datasets

MCF10A cell populations were treated with seven ligands, PBS (control), HGF, OSM, EGF, EGF+BMP2, EGF+TGFβ, and EGF+IFNG. For this paper we analyzed all but the IFNG population because initial analysis showed that it was so distinct from other cell populations that even a single marker intensity resulted in decent separability. After 48 hours, cells were fixed and subjected to cyclic immunofluorescence with 23 markers shown in Supplemental Table 1(data from the LINCS Consortium -- https://lincs.hms.harvard.edu/mcf10a/).^26^ Cells were segmented using CellPose segmentation tool,^27^ on the EGFR channel. Stains were normalized using histogram stretching to the 1^st^ and 99^th^ percentiles across intensities for the whole dataset. Image augmentations were applied for rotation, polar orientation, and size/shape (Supplemental Figure 6). Rotation is corrected by obtaining the major axis from the binary cell mask, then rotating the image using the python package OpenCV.^28^ Polar orientation was corrected by calculating the angle toward the image’s center of mass, then applying a flip/rotation to align the angle using the python Numpy package.^29^ Size/shape was corrected simultaneously by registering the cell mask to a circle target image (code available here: https://github.com/GelatinFrogs/Cells2Circles). In total, 71,314 cells were processed through this pipeline.

Bulk Reverse Phase Protein Array (RPPA) was performed by the LINCS consortium^26^ in parallel to the CYCIF imaging, on cell populations treated with the same ligands after 48 hours of exposure. The protein array incorporated 295 protein markers. As described by Akbani, R., et al.,^30^ RPPA data were median-centered and normalized by standard deviation across all samples for each component to obtain the relative protein level. The pathway score is then the sum of the relative protein level of all positive regulatory components minus that of negative regulatory components in a particular pathway. Pathway members and weights were developed through literature review. Pathways were used instead of individual proteins because the large number of proteins would decrease the significance of correlations. Despite the available bulk RPPA dataset saving a smaller sample size than the single cell CYCIF dataset, it was chosen as the secondary modality because similar ligand separation and cluster patterns were observed in both modalities, indicating an overlap in the information each contains (Supplemental Figure 7).

For correlation to CYCIF and RPPA pathways, the VAE latent space was restricted to smaller sets of aggregated features. These aggregated features were made using self-correlation of VAE features across individual cell metrics and averaging the VAE features for resulting hierarchical clusters (Fig. 4 and Supplemental Figure 4). This was done to reduce the feature dimensionality and reduce spurious correlations in the biological findings. Representative images for each cluster were done by finding cells with a high average expression for all features within the cluster. For other analyses of VAE features, ME-VAE encoding features were restricted to 18 single features for each. The dimension of 18 was chosen because it is the number of mutual markers between the RPPA and CYCIF datasets. Explanatory features were chosen from the VAE encodings such that the inter-cluster variability was maximized and the intra-cluster variability was minimized using the following equation:

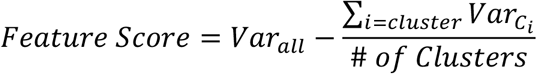

### B. VAE Models

To allow for fair comparison, the structure of the encoder and decoder blocks were kept consistent between networks, and the same latent dimension was used for all models for a given dataset (64 for the 1-channel dataset, 512 for the 23-channel dataset). All models were trained for 10 epochs on the NVIDIA P100 with 100GB of memory, but the ME-VAE architecture can work on any NVIDIA GPU. Standard VAEs with matching pairs of single cell images were used to establish baseline performance (Fig. 1c):

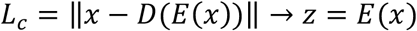

By using an image randomly transformed w.r.t. a dominant feature as the input and controlling for the uninformative feature in the output image (Fig. 1d), the model can self-supervise the transformation and will only encode novel features since the controlled features (such as rotation) no longer aid reconstruction:

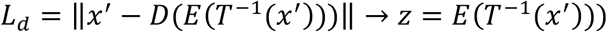

where x′ represents an image that has been transformed with a known transformation to remove one or more uninformative features and T-^1^(·) represents a transformation of the controlled image to create a dominant uninformative feature at a random degree.

The proposed multi-encoder architecture uses multiple transformed inputs with separate encoder blocks, where each block controls for a separate uninformative feature, and a single decoder block uses the shared latent space for reconstruction (Fig. 1d):

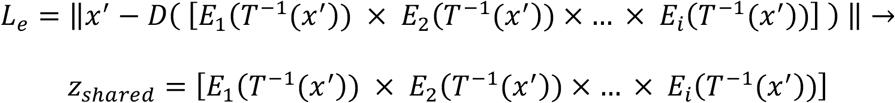

where each encoder’s individual latent space is combined in an elementwise multiplication layer and T*i*-1(·) represents a different random transformation for individual uninformative features such as rotation, polar orientation, size, shape, etc. respectively. The shared latent space of the multi-encoder forces the deep learning model to encode features that are shared between each transformation, reinforcing the shared mutual information and eliminating the non-shared transformational information.

The multi-encoder architecture allows for image pairs to be randomly transformed, which can act as a balancing agent for imbalanced features. Furthermore, the corrected outputs serve as a weakly self-supervised signal for the model. With the extra information from the additional inputs, the model is able to overcome more complex transformations that failed in the corrected output architecture in Figure 1d, when multiple corrections are attempted. Paired images also serve one additional benefit of allowing for features to be retained in parallel encoders that might be lost due to artifacts in other corrections, i.e. artifacts within a polarity correction encoder will not be present in a rotation correction encoder.

### C. Evaluations Metrics

In order to evaluate the model’s ability to separate labeled cell populations, k-means clustering was applied to the encoding spaces using sklearn.^31^ Cluster purity was then calculated by taking the percentage of the largest population for each cluster. UMAP embeddings were calculated using the UMAP python package.^32^ Biological metrics were calculated to give VAE encoding features biological grounding (Supplemental Figure 2). Circularized cells were used for calculation because it made compartmentalization of the cell more consistent and uniform. Mean intensities were calculated for inner, middle, outer, and whole cell compartments. To calculate the radial slope, the mean intensity was taken from each radius of the circularized cell, then the linear regression of the series was calculated using the scipy.stats package^33^ in python. Self-correlations between VAE features were performed using spearman correlation and clustering was done in seaborn clustermap.^34^ Representative cluster images were chosen based on high expression of the cluster’s respective VAE features. RPPA pathways activity scores, VAE features, and biological metrics were all standardized prior to analyses using the sklearn StandardScaler function^31^ in python. Correlations between RPPA pathway activities and VAE encodings and between CYCIF and VAE encodings were both calculated using the spearman correlation. To test for separability (Fig. 5), features were first tested using ANOVA, all of which proved significant. Subsequently, the post-hoc pairwise Tukey p-test was used to calculate the significance and effect size for each ligand pair. The mean p-value and effect size were reported to illustrate average separability.

## Acknowledgements

This work was supported in part by the National Cancer Institute -- U54HG008100 (Gray), U54CA209988 (Gray), U2CCA233280 (Gray), U01 CA224012, R01 CA253860 (Chang) -- U01 CA217842 and 1U01 CA253472-01A1 (Mills; Korkut; Liang) -- and the OHSU Center for Spatial Systems Biomedicine. The resources of the Exacloud high performance computing environment developed jointly by OHSU and Intel and the technical support of the OHSU Advanced Computing Center are gratefully acknowledged.

## Author contributions

Conceptualization, L.T. and YHC.; Methodology, L.T. and Y.H.C.; Software, L.T.; Formal Analysis, L.T.; RPPA Analysis, M.D. and M. L. Writing original draft, L.T.; Writing-Review and Editing, L.T. and Y.H.C.; Supervision, G.M., J.W.G., L.H. and Y.H.C.; Funding Acquisition, J.W.G.;

## Supplemental Material

### Supplemental Figures

**Supplemental Figure 1:**
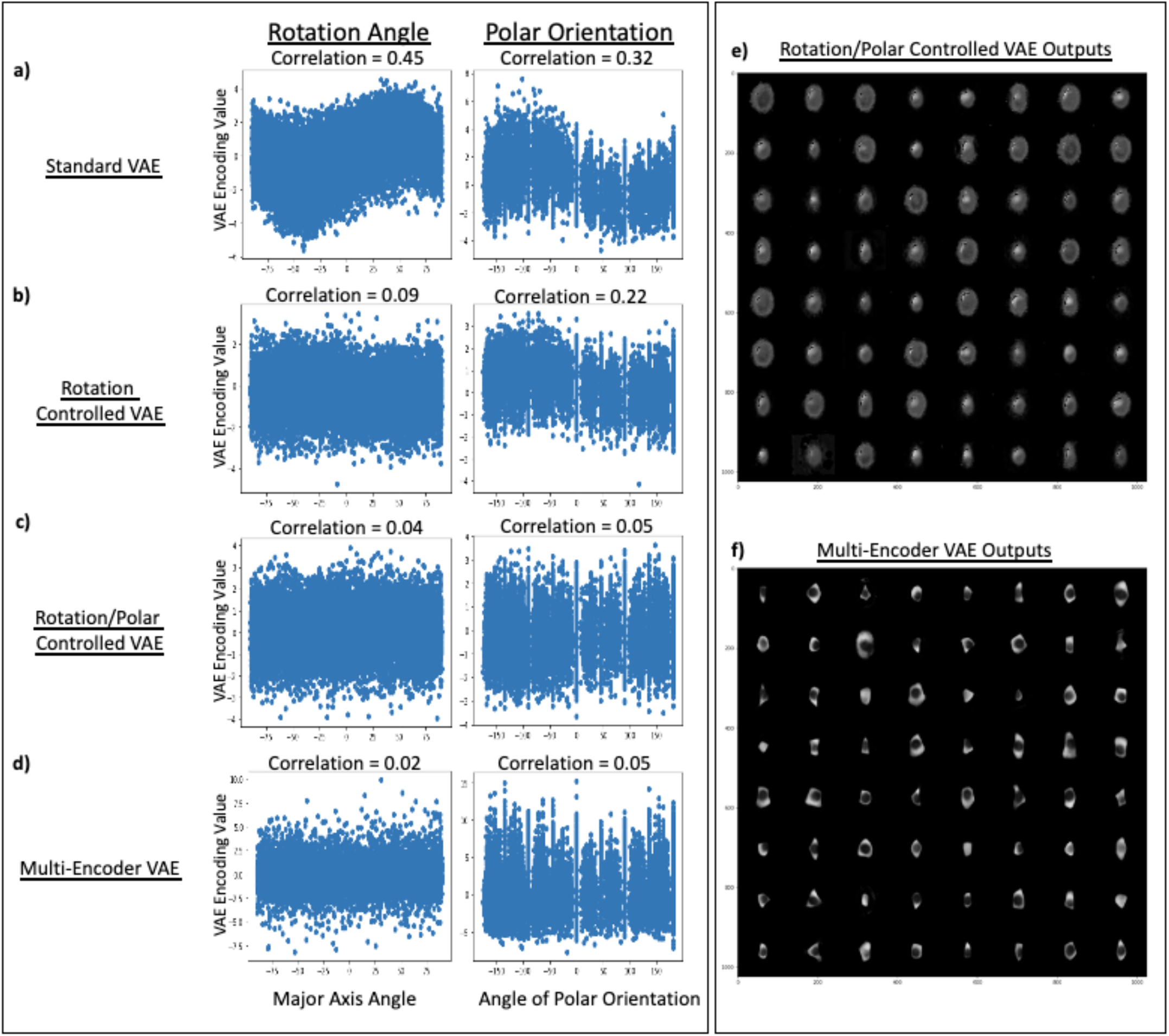
Mitigation of uninformative features for tested architectures. Encoding spaces for each VAE method were analyzed for correlation with uninformative features. Scatter plot and correlation is shown for the latent space component that had the highest correlation to given the metric. **a)** Standard VAE used as baseline to show high correlation between encoded features and undesired features. **b)** Output corrected transform invariant VAE controlling for rotation only. **c)** Output corrected transform invariant VAE controlling for rotation and polar orientation. **d)** Proposed multi-encoder VAE correcting for both rotation and polar orientation. **e**) Failed reconstruction examples from the transform invariant VAE correcting for both rotation and polar orientation. **f**) Successful recocnstruction examples from the ME-VAE correcting for rotation and polar orientation.

**Supplemental Figure 2:**
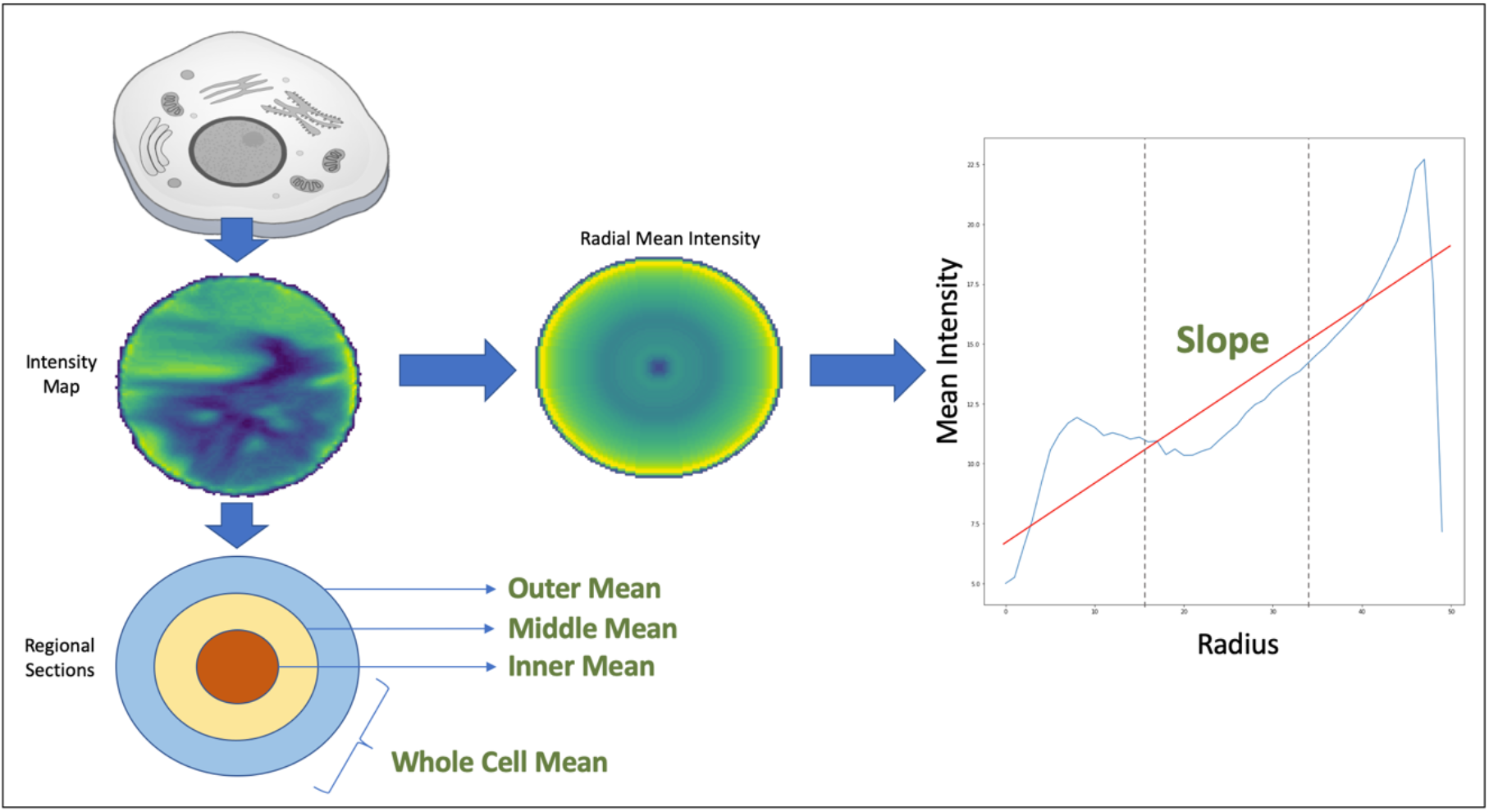
Extracted biological metrics from CYCIF. Cell intensity maps were circularized to allow easy compartmentalization. The inner, middle and outer mean intensities were extracted by dividing the cell into thirds radially. The mean intensity of the whole cell was also taken. The radial mean intensity map was created by taking the average intensity for each radius across the circularized cell. The slope of the radial mean intensity map was then taken to create a single metric for stain distribution.

**Supplemental Figure 3:**
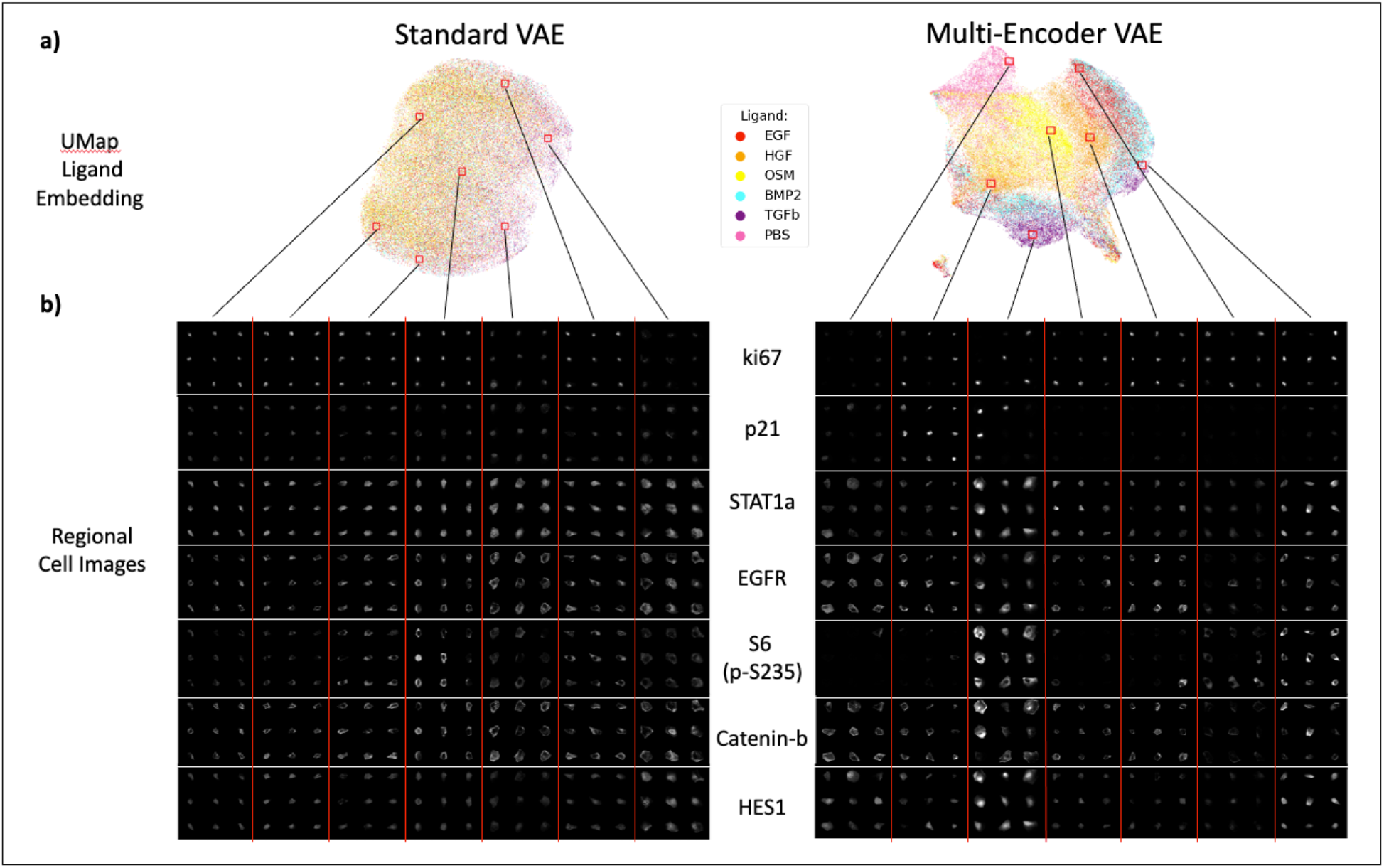
Regional cell images across UMAP clusters. Regional cell images were sampled from locations throughout UMAP space to highlight the differences in expression pattern. Stains shown were selected based on a combination of being correlated to important VAE features and hand-selection for known variance.

**Supplemental Figure 4:**
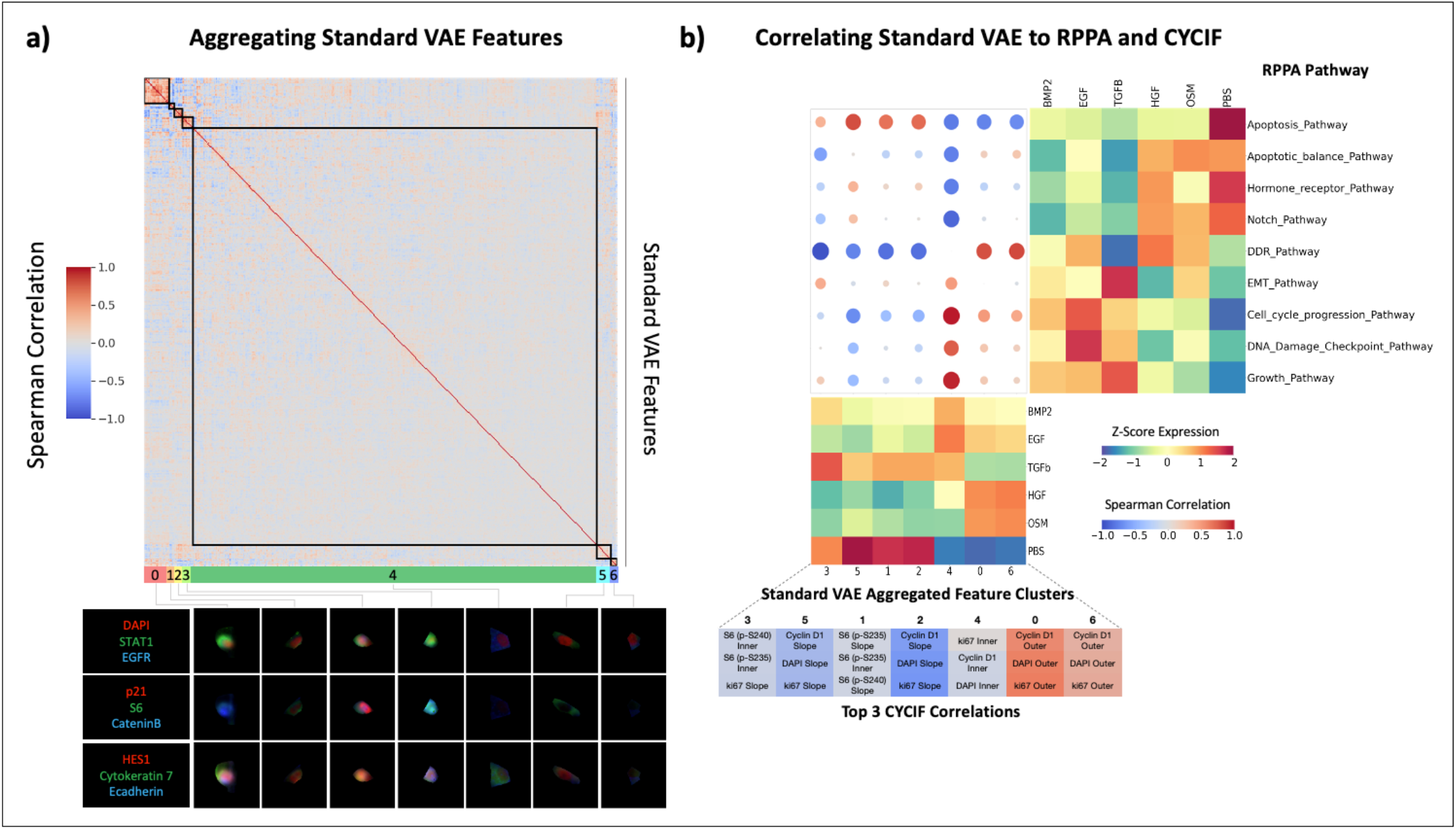
Standard-VAE feature aggregation and transitive inter-modality correlation. **a)** Using the single cell observations as features, correlations are drawn between pairs of standard VAE features. These features are then hierarchically clustered to observe patterns and reduce VAE features to aggregated feature sets. Cell images were assigned aggregated feature scores using the mean expression of each feature in a cluster. Shown are representative cells that are highly expressing for each respective cluster. **b)** Correlation matrix between RPPA pathway activity scores and standard VAE aggregated features. Samples from the two modalities were paired by their ligand treatments, resulting in a sample size of n=6. RPPA pathways and VAE features were hierarchically clustered to show prominent patterns in correlation. Standard VAE aggregated features were also correlated to several metrics of CYCIF expression (mean inner, mean middle, whole cell mean, and radial slope) for all 23 stains. This CYCIF correlation was done using the full dataset of single cell images (sample size n=73,134). The table of CYCIF correlations shows the top three correlations for each ME-VAE aggregated feature.

**Supplemental Figure 5:**
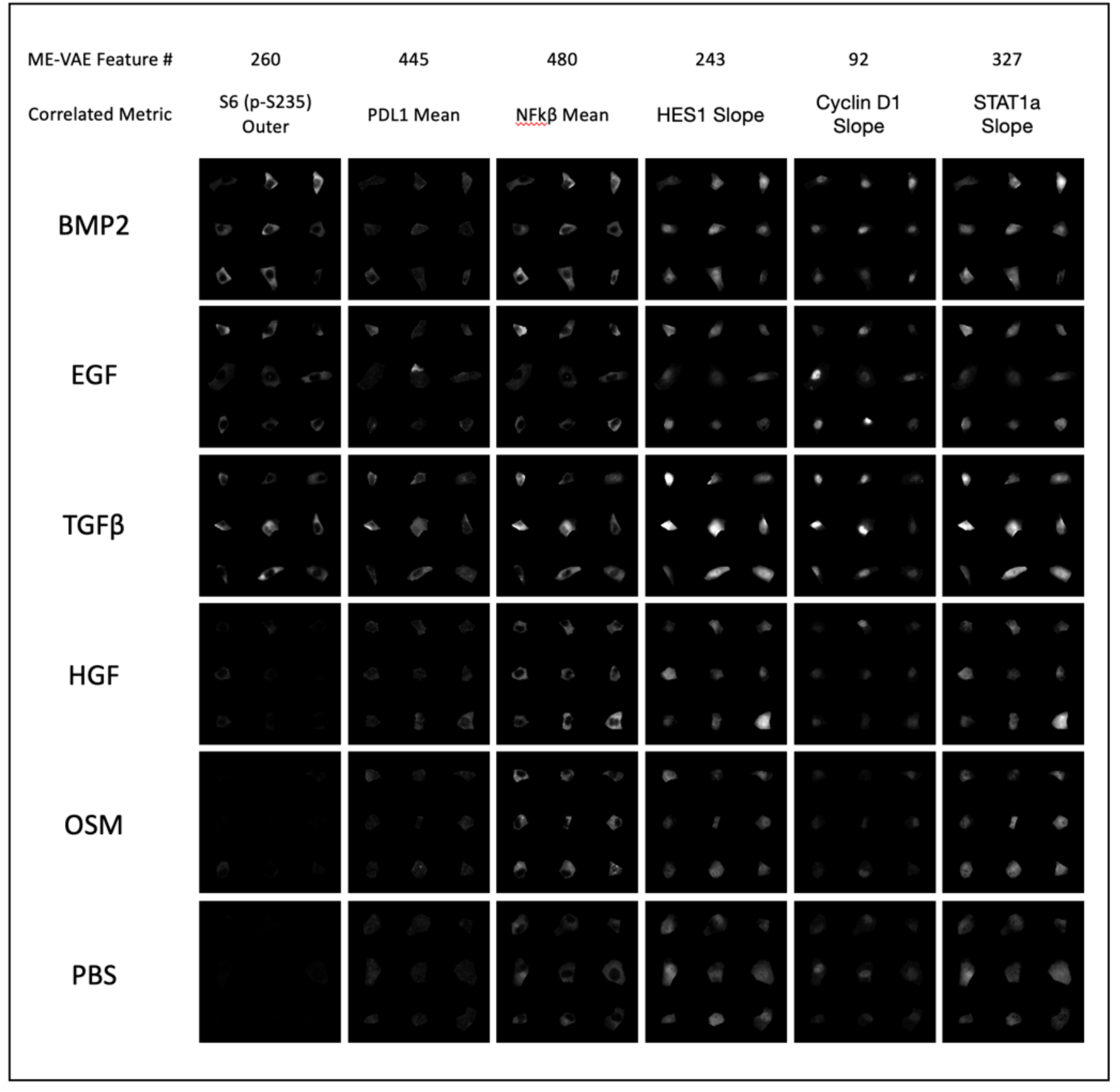
Representative cell images for each ligand treatment. Representative cell images are shown for each ligand treatment (rows) and are shown using several stains (columns). Each column also includes a VAE # that ties back to the multi-encoder feature that is highly correlated.

**Supplemental Figure 6:**
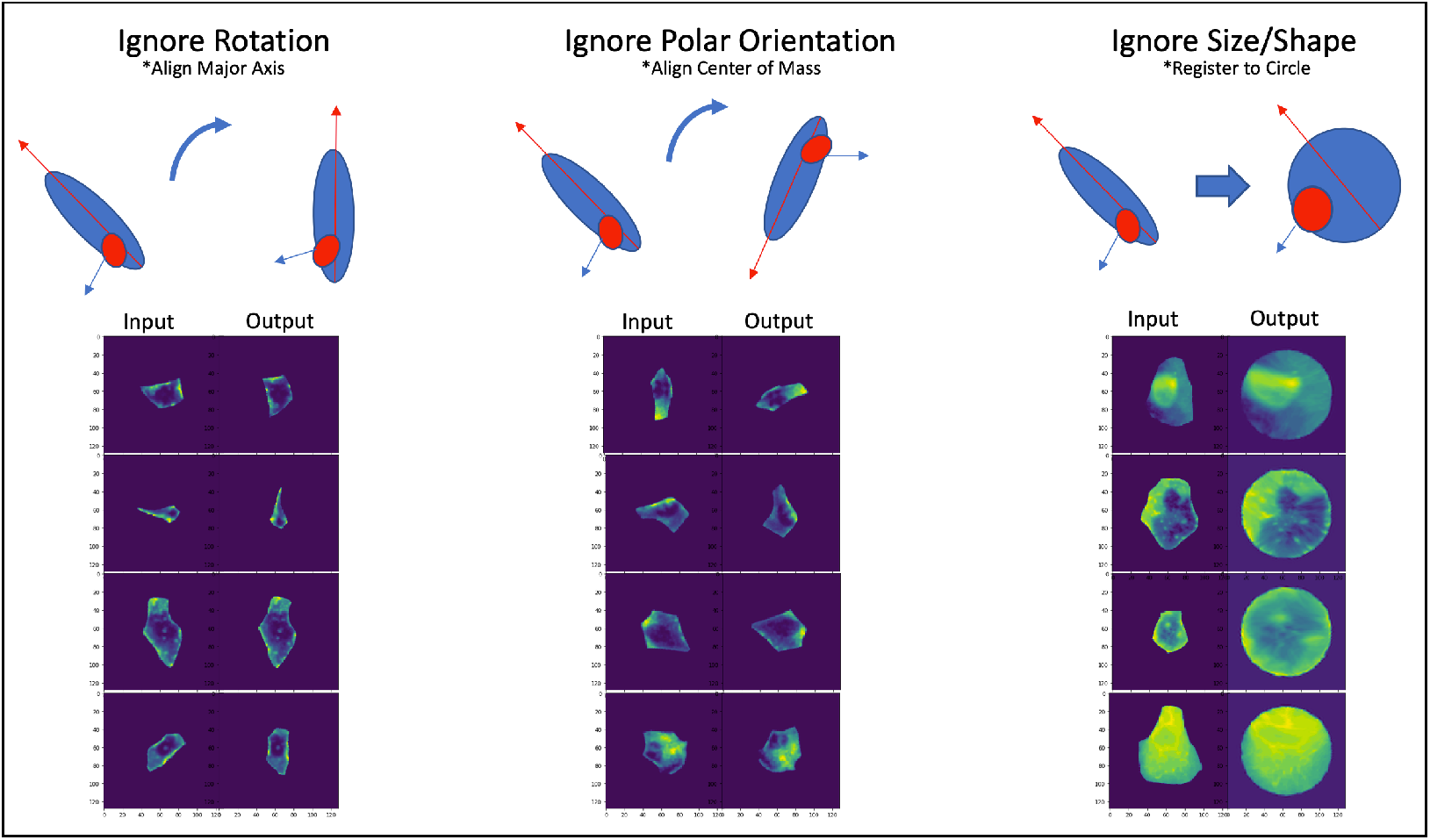
Cell image augmentation and correction. Examples of image corrections for rotation, polar orientation, and size/shape, shown using EFGR channel of randomly selected images.

**Supplemental Figure 7:**
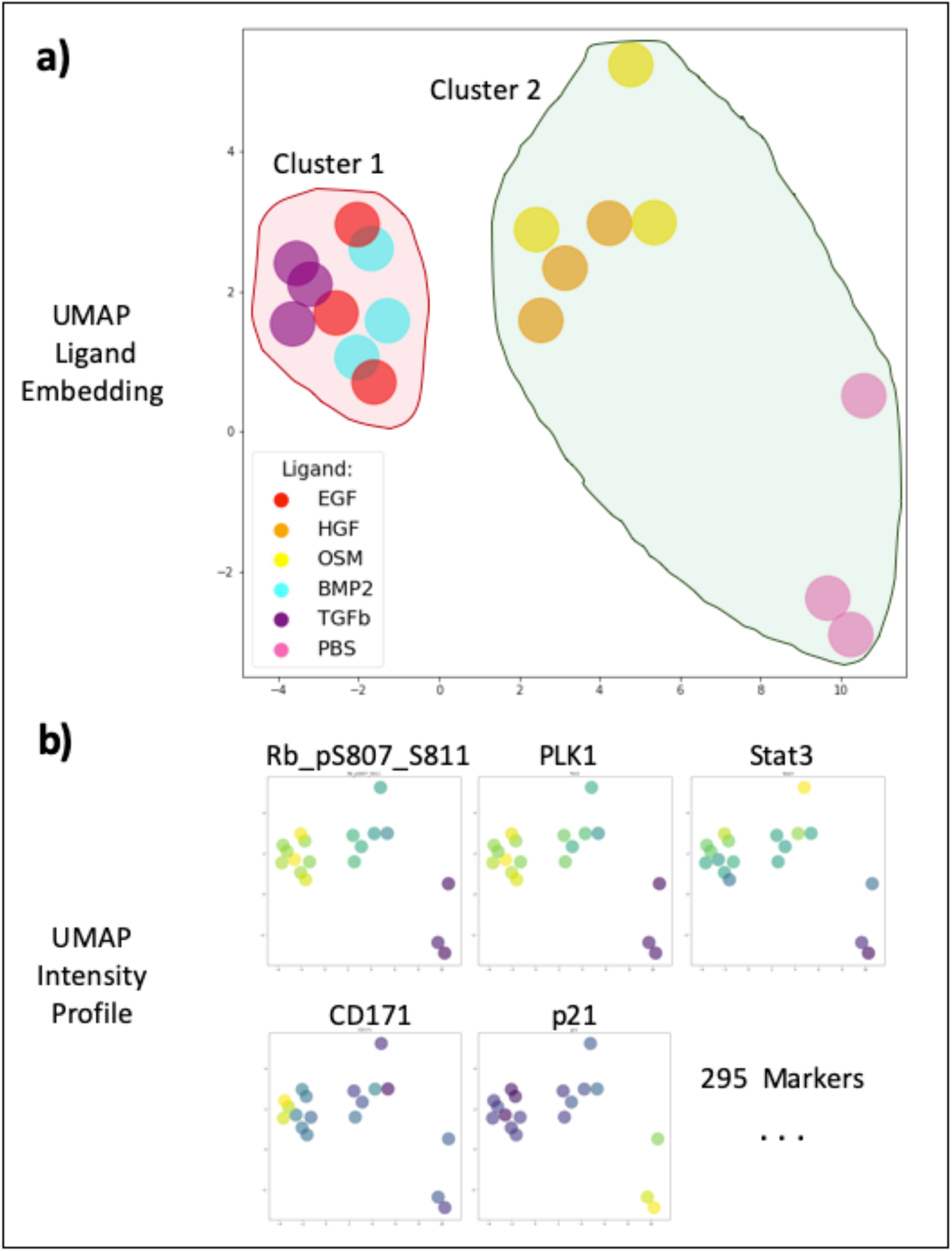
Bulk RPPA analysis and clustering. **a)** Independent analysis of the Bulk RPPA dataset shows distinct clustering of ligand populations in UMAP embeddings space where **b)** selected markers show clear patterns of distribution between the clusters.

## Supplemental Table

**Supplemental Table 1:** CYCIF Markers

| Channel | Marker |
| --- | --- |
| 1 | DAPI |
| 2 | STAT1 (p-S727) |
| 3 | Vimentin |
| 4 | Cytokeratin 7 |
| 5 | ki67 |
| 6 | S6 |
| 7 | LC3A/B |
| 8 | NFkB (p65) |
| 9 | p21 (Waf1/Cip1) |
| 10 | Catenin (Beta) |
| 11 | S6 (p-S235/S236) |
| 12 | PDL1 |
| 13 | E-cadherin |
| 14 | STAT1 (alpha-isoform) |
| 15 | HES1 |
| 16 | EGFR |
| 17 | NDG1 (p-T346) |
| 18 | STAT3 |
| 19 | S6 (p-S240/244) |
| 20 | MET |
| 21 | Cytokeratin 18 |
| 22 | Cyclin D1 |
| 23 | c-Jun |

